# Deconvolving mutation and selection reveals stage-specific drivers of thyroid cancer

**DOI:** 10.1101/2025.10.08.681166

**Authors:** Moein Rajaei, Andrew Ju, Adebowale J. Adeniran, Jeffrey P. Townsend

## Abstract

**Objective:** Mutational profiles of primary and metastatic thyroid cancer (THCA) have been examined by comparison of somatic mutation prevalences in the two stages, using *P* values to identify significant differences. However, prevalences and *P* values for mutations do not directly quantify the cancer effects of somatic variants.

**Methods:** We calculated cancer effect sizes accounting for substantial gene- and site-specific mutation rates, to quantify somatic selection across stages. This approach provides a direct measure of cancer-driving impact of new mutations and reveals the genes that drive tumorigenesis and progression, and whether mutations have greater effects in primary or metastatic THCA.

**Results:** Trinucleotide mutation profiles were similar between primary and metastatic THCA. Most canonical driver genes (*BRAF, NRAS, TP53, ATM, EIF1AX, KMT2C, NF1, RBM10, ARID1A, PIK3CA*, and *NKX2-1*) exhibited stronger selection during initial tumorigenesis (from organogenesis to primary THCA) than during progression (from primary to metastatic THCA). Notably, *TERT* mutations have been shown to be at higher prevalence in metastatic tumors, yet their strongest selection occurred earlier, during tumor initiation. In contrast, *RET* mutations exhibited the opposite trajectory, experiencing weaker selection during tumorigenesis but stronger selection during metastatic progression.

**Conclusion:** Cancer effect size analysis revealed dynamic shifts in selective pressures across THCA evolution, distinguishing genes that drive initiation from those that promote metastatic progression. This evolutionary framework provides a quantitative basis for understanding THCA pathogenesis and provides guidance for stage-specific precision-medicine therapeutic strategies.

## Background

Thyroid cancer (THCA) has one of the highest contemporary 5-year survival rates among cancers (98%) ^1^. Nevertheless,it accounted for an estimated 821,000 new cases and 44,000 deaths worldwide in 2022^2^, ranking as the seventh-most commonly diagnosed malignancy. Prognosis declines sharply with stage, from 99.8% survival for localized tumors to 97% with regional metastases, and 57.3% with distant metastases ^3^, demonstrating the importance of early diagnosis and treatment to prevent metastatic spread and improve outcomes.

Given the crucial value of understanding the differences between primary and metastatic THCA, prior studies have characterized variant profiles of tumors by reporting differences in prevalence between stages, accompanied by their statistical significance ^4^, However, prevalence conveys only the proportion of patients carrying a variant, and *P* values convey only the probability of observed differences given the data. They are related to, but are not measures of oncogenic effect ^5^. A more comprehensive understanding therefore requires quantification of mutational patterns, background mutation rates, and selective pressures on driver genes across disease stages.

To guide basic and translational research, we analyzed 2,145 primary and 1,374 metastatic THCA sequences from Maximo *et al*^*4*^, Masoodi *et al*^*6*^, TCGA, and GENIE. Quantifying gene- and site-specific mutation rates, we established neutral expectations, separating mutational processes from selective pressures. We then quantified somatic selection on driver genes, distinguishing those most crucial to tumor initiation from those driving metastatic progression.

## Methods

Of 475 target-sequenced primary and metastatic TCs reported by Maximo *et al*. ^4^, we analyzed the 458 non-follicular tumors for which non-synonymous single-nucleotide variants (SNVs) were provided. We also included whole-exome sequences from TCGA, targeted panel data from GENIE Release 15.1 ^7^, and 14 matched primary-metastasis pairs from Masoodi *et al*. ^6^. In total, our dataset comprised 2,145 primary and 1,374 metastatic tumors, aligned to the human reference genome Hg19. TCGA data originally aligned to Hg38 was converted to Hg19 using Liftover.

We analyzed the trinucleotide mutational profiles of primary and metastatic THCA using MutationalPatterns v3.8.1 ^8^ and cancereffectsizeR v2.9.0 ^9^. Following Cannataro et al. ^10^, we excluded mutational signatures previously determined to be absent in THCA ^11^. Trinucleotide mutation rates were convolved with gene-specific background rates ^12^ to estimate neutral expectations for each site, and the sequential model ^9^ was applied to quantify somatic selection during both primary tumorigenesis and metastatic progression. Analyses focused on the sixteen most prevalent thyroid cancer genes reported by Maximo et al. ^4^. Variants within genes were modeled with site-specific mutation rates but as sharing a single invariant cancer effect size. This approach enabled quantification of stage-specific differences in somatic selection intensity, providing insight into the mutations driving thyroid cancer initiation and progression and informing the basis for stage-targeted precision therapies.

## Results

The overall trinucleotide mutational landscape was largely similar between primary and metastatic THCA. However, GTG → GAG, TTG → TCG, and ATG → ACG substitutions were at substantially higher proportions in metastases. These differences in trinucleotide mutation profile were insufficient to cause the differences in variant prevalence between primary and metastatic THCA.

Neutral mutation accumulation was significantly higher in metastatic tumors than in primary tumors (rank-biserial correlation = 0.63; *P* < 2.2 × 10^−16^; Mann–Whitney U test), and across all 16 genes analyzed, neutral mutation rates were elevated in metastatic disease (**Table 1**). Mutations of *RET* were more prevalent in metastatic tumors than in primary tumors (**Fig. 1A**). Consistent with this pattern of prevalence, our analysis showed that *RET* mutations were subject to weaker selection during primary tumorigenesis, and stronger selection during metastatic progression (**Fig. 1B**). In contrast, *NRAS* and *KMT2C* mutations exhibited the opposite trajectory: they were much more strongly selected during primary tumorigenesis than during progression to metastasis, even though their prevalence was similar across stages.

**Table 1.** Gene-level mutation rates in primary and metastatic thyroid tumors.

| Gene | Gene mutation rate |  |
| --- | --- | --- |
|  | Organogenesis to primary tumor | Primary tumor to metastasis |
| *PIK3CA* | $8.4 \times 10^{-8}$ | $1.3 \times 10^{-7}$ |
| *BCOR* | $1.1 \times 10^{-7}$ | $1.7 \times 10^{-7}$ |
| *ATM* | $7.4 \times 10^{-8}$ | $1.8 \times 10^{-7}$ |
| *ARID1A* | $1.1 \times 10^{-7}$ | $1.9 \times 10^{-7}$ |
| *EIF1AX* | $1.1 \times 10^{-7}$ | $2.1 \times 10^{-7}$ |
| *KMT2C* | $7.7 \times 10^{-8}$ | $2.2 \times 10^{-7}$ |
| *RET* | $1.7 \times 10^{-7}$ | $2.3 \times 10^{-7}$ |
| *NF1* | $9.4 \times 10^{-8}$ | $2.5 \times 10^{-7}$ |
| *BRAF* | $8.4 \times 10^{-8}$ | $2.5 \times 10^{-7}$ |
| *NRAS* | $1.2 \times 10^{-7}$ | $2.5 \times 10^{-7}$ |
| *RBM10* | $1.0 \times 10^{-7}$ | $2.9 \times 10^{-7}$ |
| *TP53* | $1.3 \times 10^{-7}$ | $3.0 \times 10^{-7}$ |
| *CDKN2B* | $1.7 \times 10^{-7}$ | $4.1 \times 10^{-7}$ |
| *TERT* | $1.5 \times 10^{-7}$ | $4.2 \times 10^{-7}$ |
| *TBX3* | $1.7 \times 10^{-7}$ | $4.7 \times 10^{-7}$ |
| *NKX2-1* | $1.8 \times 10^{-7}$ | $1.1 \times 10^{-6}$ |

**Figure 1.**
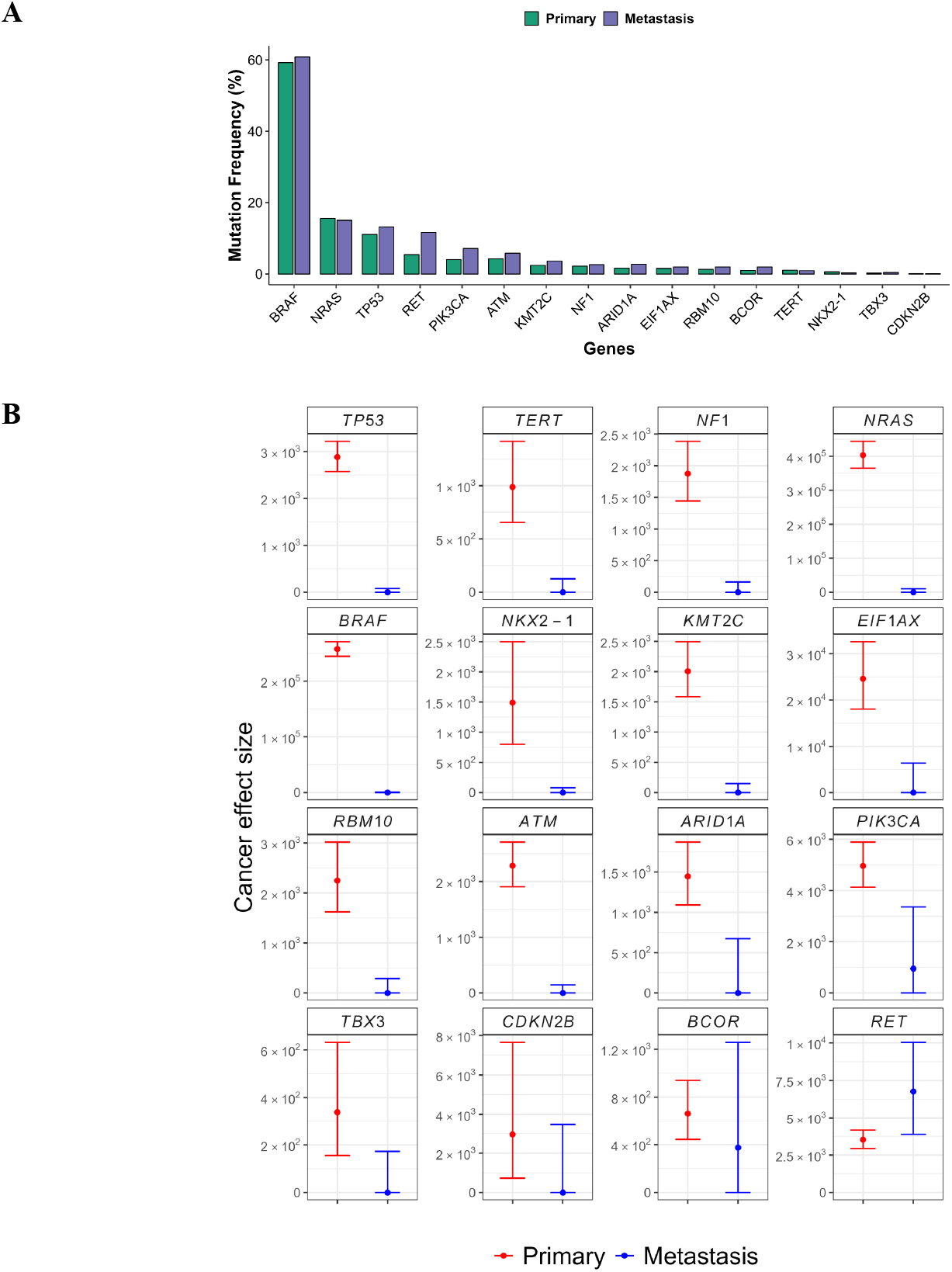
(**A**) Prevalence of mutations in 16 selected genes across primary and metastatic thyroid cancers. (**B**) Cancer effect sizes of somatic variants in oncogenic sites / regions / domains of 16 genes that are known to act as drivers in thyroid cancer tumorigenesis and metastasis.

Metastases carried *BRAF* mutations at a frequency comparable to primary tumors, and *BRAF* mutation rates were increasing but fairly consistent between stages (2.5 × 10^−7^ *vs*. 8.4 × 10^−8^). In contrast new *BRAF* mutations were subject to much stronger selection during the transition from organogenesis to primary THCA than during progression to metastasis (**Fig. 1B)**.

Just as a similar mutation frequency across stages does not reliably scale with selective pressure, a higher frequency of gene mutation in a later stage does not necessarily equate to higher selection. The prevalence of *TP53* mutations was higher in metastatic tumors than in primary tumors. However, the selection intensity for *TP53* mutations occurring during progression to metastasis was lower than it was for mutations occurring during primary tumorigenesis (**Fig. 1B)**. In this case, variant prevalence increase and strength of selection on the variant were decoupled, with prevalence increasing even as selection intensity decreased.

Mutations in *TERT, ATM, NF1, NKX2-1, EIF1AX, RBM10, PIK3CA*, and *ARID1A* were subject to stronger somatic selection from organogenesis through primary tumorigenesis than during progression to metastasis (**Fig. 1B**). Mutations in *TBX3, CDKN2B*, and *BCOR* genes did not exhibit statistically significant differences in the intensity of selection between primary tumorigenesis and progression to metastasis. Overall, most driver genes demonstrated higher selective pressure earlier in disease progression, with only a subset maintaining similar intensities across stages.

## Discussion

Our analysis demonstrated that *TP53* and *RET* mutations both exhibited higher prevalence in metastatic than in primary tumors. However, their selective dynamics were divergent: novel *TP53* mutations were under weaker selection during progression to metastasis than during primary tumorigenesis, whereas *RET* mutations were under significantly stronger selection during metastatic progression—consistent with a prior study demonstrating a functional role of *RET* in metastatic THCA ^16^. These findings indicate that therapeutic inhibition of *RET* might be beneficial even in tumors that have not yet acquired *RET* mutations, as such intervention could potentially forestall metastatic progression. By accounting for underlying neutral mutation rates and quantifying cancer effect, these analyses demonstrate the enhanced power and accuracy obtained when deconvolving mutation and selection differences across stages of cancer.

Our analysis classified tumors as “primary” or “metastatic”, a relevant but coarse binary discretization of the continuum of thyroid cancer progression. Such discretization obscures intermediate states such as locally advanced or regionally metastatic disease, which may exert specific selective dynamics on novel gene variants. Nonetheless, the contrast between localized primaries and established distant metastases maximizes power with a limited sample size and provides a clear and biologically meaningful comparison. This dichotomy enables robust detection of shifts in selective pressure, while also motivating future work that dissects finer-grained transitions.

Our analysis relied on bulk sequencing, which integrates signals from genetically heterogeneous tumor cell populations and does not distinguish clonal from subclonal mutations or model frequency-dependent selection among competing variants. Higher-resolution approaches such as single-cell sequencing or spatial genomics will be needed to capture these dynamics of clonal competition and dissemination. Nevertheless, this limitation does not compromise the accuracy of our estimates of population-level selective pressures, which reflect longer-term molecular evolutionary outcomes determining frequencies of variants across tumors.

Incorporation of gene- and site-specific mutation rates to calculate cancer effect enables quantification of somatic selection and the contributions of individual variants and genes to thyroid tumorigenesis and progression. However, this approach does not elucidate the underlying biological mechanisms that drive that selection. It identifies which mutations are favored across disease stages, but cannot reveal whether selection arises from altered signaling pathways, changes in the tumor microenvironment, or other molecular and cellular processes. Accordingly, these findings should motivate functional studies, pathway analyses, and integrative genomic approaches that determine why specific variants confer selective advantages. Such complementary research will not only validate the drivers identified here but also provide mechanistic insights that can guide the development of effective, stage-specific therapeutic strategies.

Finally, our analysis was limited to coding single-nucleotide variants, whereas other processes—copy number alterations, gene fusions, epigenetic modifications, transcriptional reprogramming, and interactions with the tumor microenvironment—also play critical roles in metastatic progression. Thus, our framework captures only part of the complex landscape of thyroid cancer evolution. Extending cancer effect size analyses to encompass these additional molecular layers will be an important next step, enabling a more comprehensive view of the selective forces driving thyroid cancer initiation and metastatic spread.

## Acknowledgements

The authors would like to acknowledge the American Association for Cancer Research for the development of the AACR Project GENIE registry, as well as members of the consortium for their commitment to data sharing. Interpretations are the responsibility of study authors.

